# Isolation and Characterization of a *Rhizobium* Bacterium Associated with the Toxic Dinoflagellate *Gambierdiscus balechii*

**DOI:** 10.1101/789107

**Authors:** Zhen Wu, Xiaohong Yang, Senjie Lin, Wai Hin Lee, Paul K.S. Lam

## Abstract

Algae-bacteria associations are increasingly being recognized to be important in shaping the growth of both algae and bacteria. Bacteria belonging to order Rhizobiales are important symbionts of legumes often developing as nodules on plant roots, but have not been widely documented in association with algae. Here, we detected, isolated, and characterized a *Rhizobium* species from the toxic benthic dinoflagellate *Gambierdiscus* culture. The sequence of 16S rDNA showed 99% identity with that of *Rhizobium rosettiformans*. To further characterize the bacterium, we amplified and sequenced a cell wall hydrolase (CWH)-encoding gene; phylogenetic analysis indicated that this sequence was similar to the homologs of *Martellela* sp. and *Hoeflea* sp, of order Rhizobiales. We performed PCR using *nif*H primers to determine whether this bacterium can fix N_2_; however, the results of sequencing analysis showed that it was closer to chlorophyllide *a* reductase-encoding gene (*bch*X), which is similar to *nif*H. Results of 16S rDNA qPCR showed that compared to that in the early exponential phase, the abundance of this bacterium increased during the late exponential growth phase of *Gambierdiscus*. When the dinoflagellate culture was subjected to N limitation, the abundance of the bacterium represented by both 16S rDNA and CWH increased. Based on these results and published literature, it is apparent that this *Rhizobium* bacterium benefits from the association with *Gambierdiscus* by hydrolyzing and utilizing the extracellular organic matter exudates released by the dinoflagellate. This is the first report of *Rhizobium* species being associated with dinoflagellates, which will shed light on the algae-bacteria relationships.

**IMPORTANCE:** Phytoplankton are the undisputed primary producers in the aquatic ecosystems and contribute approximately half of the global net primary productivity.

Dinoflagellates are one of the most important phytoplankton in the marine ecosystems. Commonly, they do not exist autonomously in the marine environment but rather co-live with many bacteria that interact with dinoflagellates, producing a dynamic microbial ecosystem. Their interactions play a major role in important processes such as carbon fluxes and nutrient regeneration in the ocean, ultimately influencing the global carbon cycle and the climate. Hence, there is a need to understand the association and relationships between dinoflagellates and bacteria. Here, we tried to elucidate these interactions through isolating and characterizing a bacterium from a benthic toxic dinoflagellate culture. Our study is the first report of such bacterium being recorded to be associated with a dinoflagellate in this genus, providing new insights into the dinoflagellate-bacteria association for future research.

## INTRODUCTION

Algae-bacteria associations have increasingly been recognized to be important in shaping the growth of both algae and bacteria [1]. Many algae rely on vitamins produced by the bacterial community [2–7]. The symbiotic relationships between algae and bacteria can also provide algae with major nutrients such as phosphorus, nitrogen (N) [8–10], and growth regulators [11–13]. Furthermore, bacteria can be a potential food source for the mixotrophic algae [14]. In return, algae can provide organic carbon as well as nutrients for bacteria, representing a microhabitat for these associated microbes [15, 16].

Among the diverse types of algae-associated bacteria documented so far, *Roseobacter* [17], *Flavobacteria* [18], and Proteobacteria [19] are the most common. These bacteria form a mutualistic [3] or pathogenic relationship [20] with the host algae. The bacteria reported to associate with dinoflagellates include *Dinoroseobacter* spp. [21] and *Marinobacter* spp. [22]. Bacteria of order Rhizobiales are important symbionts of legumes often developing as nodules on plant roots, where these bacteria fix N_2_ and support plant growth [23]. However, the association of this type of bacteria with algae has not been widely documented, and so far, its association has been mainly limited to green algae [24] and brown algae [25]. Some species in this group of bacteria have been reported to perform photosynthesis as they contain the chlorophyllide *a* reductase, which is involved in the synthesis of bacteriochlorophyll [26] and shares amino acid sequence identity with the nitrogenase iron protein, a key enzyme in N_2_ fixation [27].

*Gambierdiscus balechii* is a benthic dinoflagellate that can produce ciguatoxins and had induced ciguatera fish poisoning in the Republic of Kiribati [28]. As benthic epiphytic dinoflagellates, *Gambierdiscus* spp. are often found in association with macro-algae in environments where nutrients are low and growth can be limited in the water column [29]. Although *Gambierdiscus* spp. survive as photoautotrophs, they can still use the exogenous organic carbon substrates for growth under light and dark conditions [30], suggesting the potential mixotrophic capacity of this genus. In this study, we aimed to identify and characterize the bacterium that contains the cell wall hydrolase (CWH)-encoding gene, which theoretically might impact the interaction between the bacterium and the host dinoflagellate. In particular, we detected, isolated, and characterized a *Rhizobium* species harboring CWH from the toxic benthic dinoflagellate *Gambierdiscus* culture. We set up experiments using various nitrogen concentrations and growth phases to determine whether and how the bacterium can benefit from *G. balechii* or whether it supports algal growth by supplying nutrients. The results were used to test two hypotheses:1) both *G. balechii* and the associated bacterium benefit from the co-growth in the culture; 2) the bacterium harbors the nitrogenase iron protein gene (*nif*H) and is potentially capable of fixing N_2_ to support algal growth. The verification of these two hypotheses will shed light on the symbiotic relationship between dinoflagellates and bacteria.

## RESULTS

### Phylogenies of 16S rDNA and CWH gene sequences from the isolated bacterium

*G. balechii* exhibited exponential growth when a sample was collected for bacterium isolation. Random clones were selected when colonies appeared on the marine agar plate. CWH primers were used to screen the bacterial clones that contained the CWH gene. Out of these selected clones, 50 % yielded positive PCR results.

Sequencing of 16S rDNA amplicons from the CWH-positive colonies revealed *Bacterium* spp., *Bacillus* spp., *Rhizibium* spp., and *Flavobacterium* spp. to be the closest matches. A *Rhizobium* species was selected for further characterization. The sequence of the isolated bacterium species (1,362 bp) was 99.93% identical to that of *Rhizobium rosettiformans* strain W3, which was isolated from groundwater from Lucknow, India [31]. Based on the commonly used species-delineating criterion of 16S rDNA identity (97% for prokaryotes), this bacterial strain was classified as *Rhizobium rosettiformans* strain GAMBA-01. The result of phylogenetic analysis verified that this bacterial strain belonged to the same sub-cluster as other *Rhizobium* strains (Fig. 1).

**FIG. 1.**
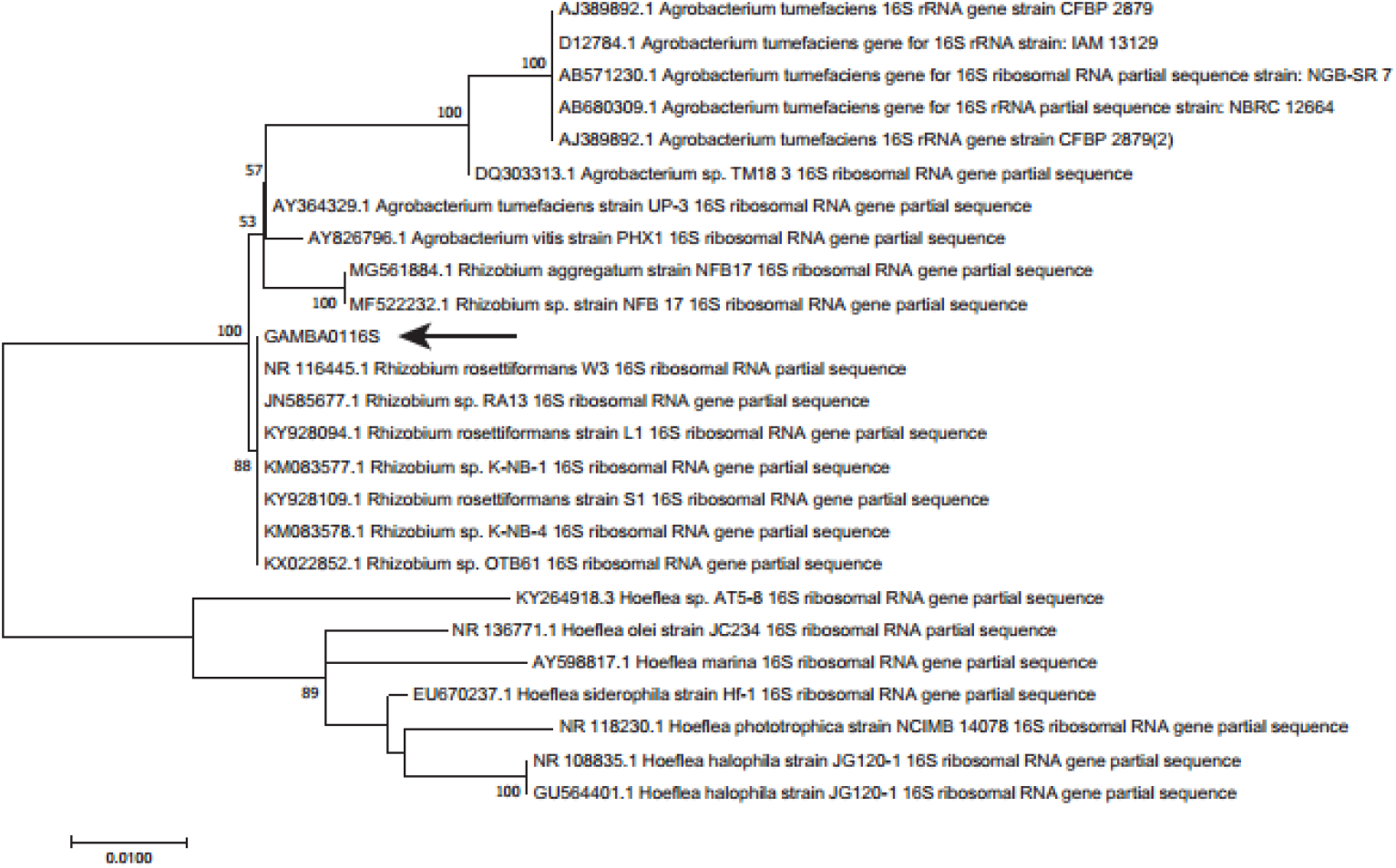
Maximum likelihood tree based on 16S rDNA sequences showing the phylogenetic position of the bacterium isolated from *G. balechii*. The sequence from our strain is marked by an arrow. Values at nodes are bootstrap support values (only those > 50% are shown).

Sequencing and phylogenetic analyses of CWH from the isolated *R. rosettiformans* strain GAMBA-01 showed that it was affiliated to *Hoeflea* sp., *Martelella* sp., and *R. flavum*, which are all genera in the order Rhizobiales (Fig. 2.). Conserved domain searches in BLASTp revealed that the CWH sequence contained a conserved domain of *cwl*J, which has been reported to be involved in spore germination [32, 33].

**FIG. 2.**
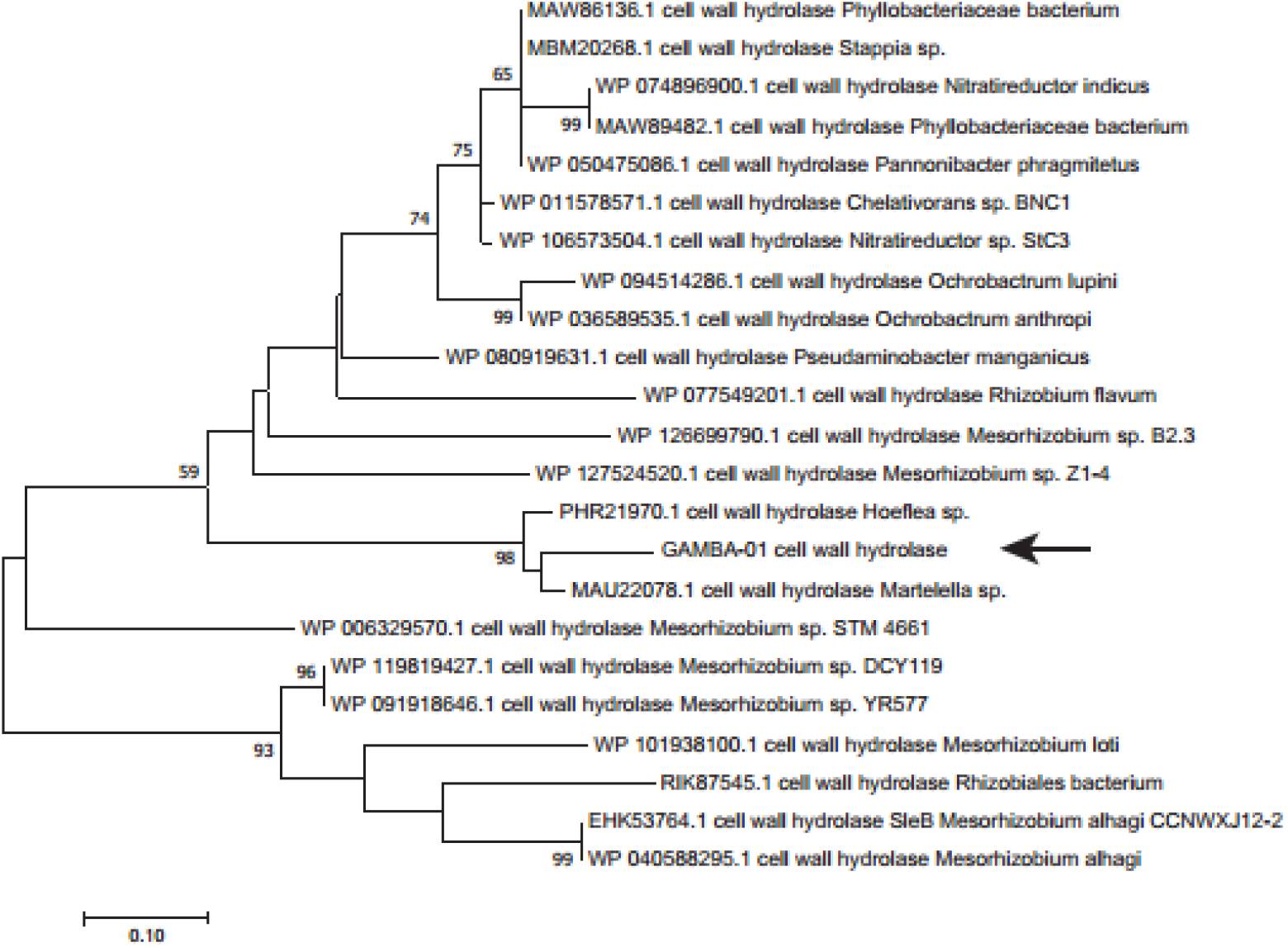
Maximum likeliood tree based on CWH amino acid sequences from bacterial strains showing the phylogenetic position of the CWH gene from the bacterium. The sequence from our strain is marked by an arrow. Values at nodes are bootstrap support values (only those > 50% are shown).

### Isolation of *bch*X using *nif*H PCR

The *nif*H gene primers Nh21F and nifH3 were used to amplify the *nif*H gene or its homologs. However, *nif*H was not detected in the amplicon after sequencing. Instead, the 593-bp PCR product was most similar (97.08% identity) to *bch*X, encoding the chlorophyllide *a* reductase iron protein subunit X of Rhizobiales bacteria. *bch*X is a homolog of *nif*H (Fig. 3) and functions in the bacteriochlorophyll *a* biosynthetic process. The deduced amino acid sequence of BchX (197 residues) from *R. rosettiformans* strain GAMBA-01 was 39.5% identical to that of NifH from *R. rosettiformans* strain W3. This isolated BchX contains a conserved domain of 4Fe-4S iron sulfur cluster binding protein that also belongs to the NifH/FrxC family. Alignments of the isolated gene product with NifH or NifH homologs showed that the conserved residues of the 4Fe-4S iron sulfur cluster-ligating cysteines are invariant, suggesting that these enzymes might function similarly (Fig. 3).

**FIG. 3.**
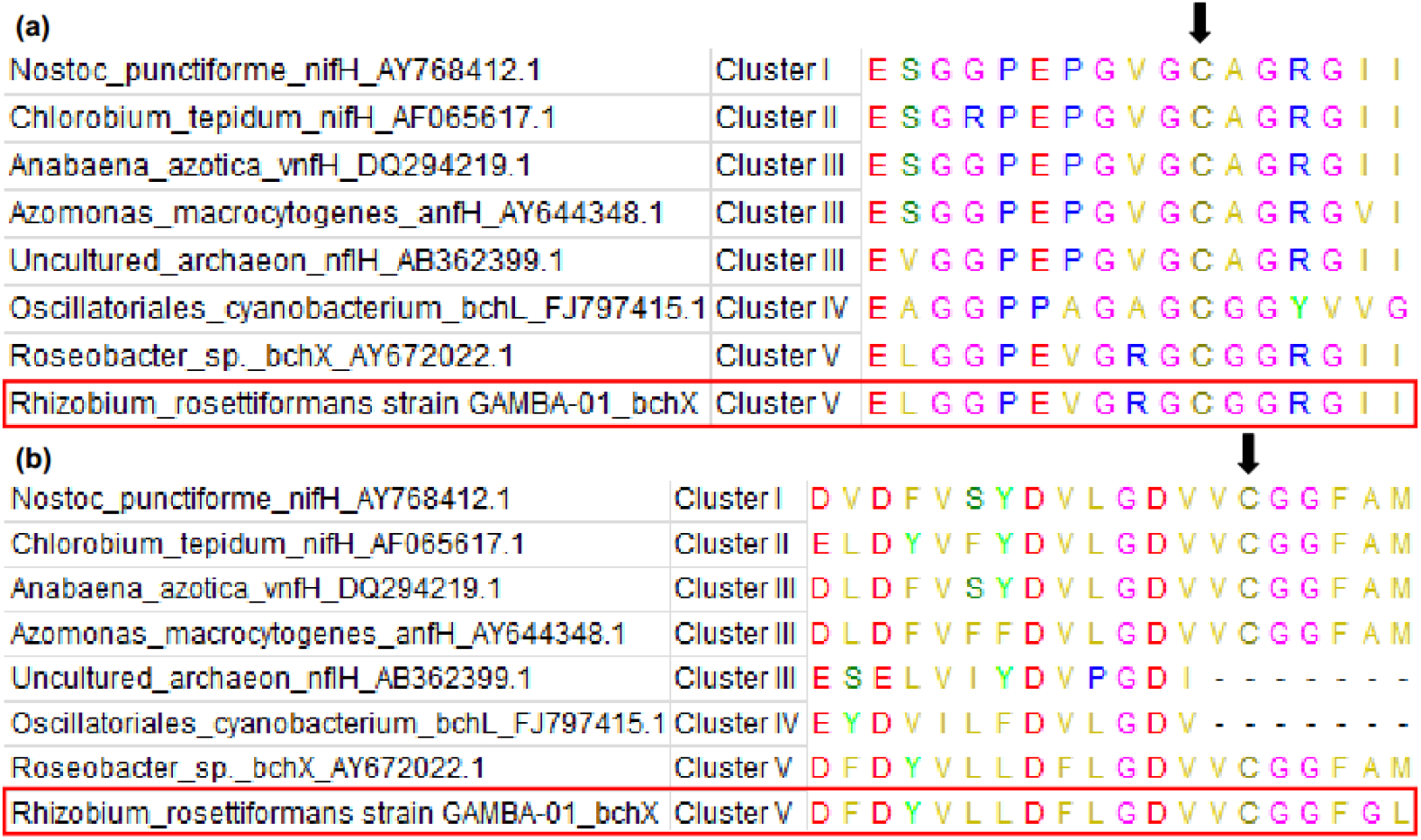
Alignment of residues surrounding the 4Fe-4S coordinating cysteine residues (vertical arrows) for NifH and NifH homologs. The sequence from our strain is boxed. (a) Cys97 of the 4Fe-4S cluster ligand; (b) Cys132 of the 4Fe-4S cluster ligand.

### Effects of N limitation on the growth of *G. balechii*-associated *R. rosettiformans*

Starting with the same *G. balechii* cell concentration, the dinoflagellate in both the N-replete group and N-limited group grew similarly until 34 days, after which their growth curves started to diverge (Fig. 4). This indicated that *G. balechii* entered N-limited physiological conditions after 34 days. The cells of the N-replete group maintained exponential growth after the 10^th^ day with the maximum growth rate of 0.22 d^−1^ (Fig. 4). The maximum cell concentration was 3,015 cells mL^−1^ on the 61-day experimental period (Fig. 4). In contrast, the maximum cell concentration for the N-limited culture was 557 cells mL^−1^ and the maximum growth rate was 0.11 d^−1^ (Fig. 4).

**FIG. 4.**
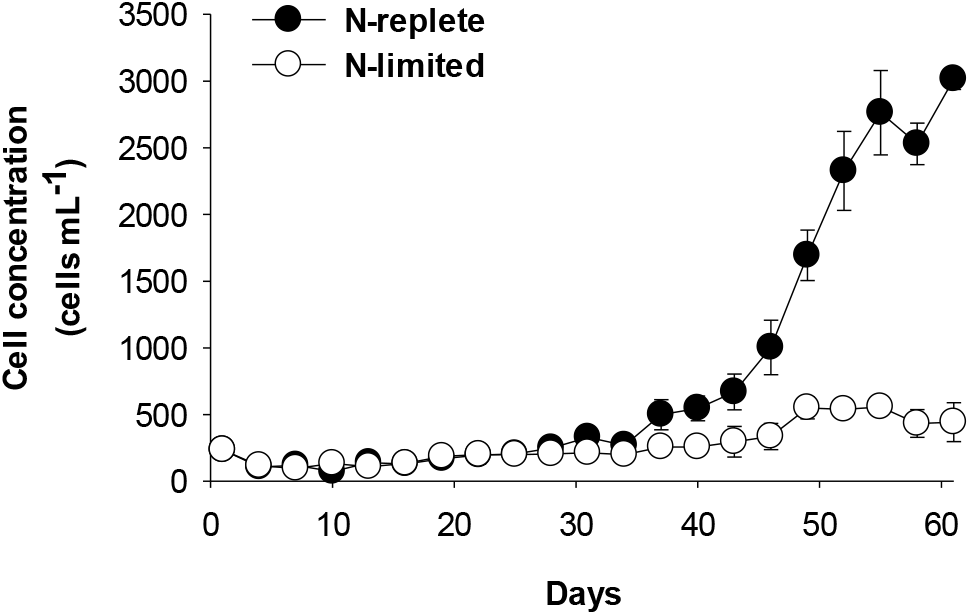
Cell concentration of *G. balechii* under N-replete and N-limited conditions.

The chlorophyll *a* contents of *G. balechii* cells from both N-replete and N-limited groups increased initially but decreased overtime after reaching the peak value (Fig. 5a). In general, chlorophyll *a* content per cell was higher in the N-replete group than in the N-limited group (Fig. 5a). The highest chlorophyll *a* value was 0.28 ng cell^−1^ on day 25 in the N-replete group and 0.21 ng cell^−1^ in the N-limited group on day 31. Chlorophyll *a* content in N-limited cultures declined rapidly after day 31, and was significantly lower than that in the N-replete group (P < 0.05) (Fig. 5a). Cell sizes were similar from day 40 to day 49 in the N-replete and N-limited groups, but were higher in the N-limited group after 49 days with significant difference on day 52 and day 55 (P < 0.05) (Fig. 5b).

**FIG. 5.**
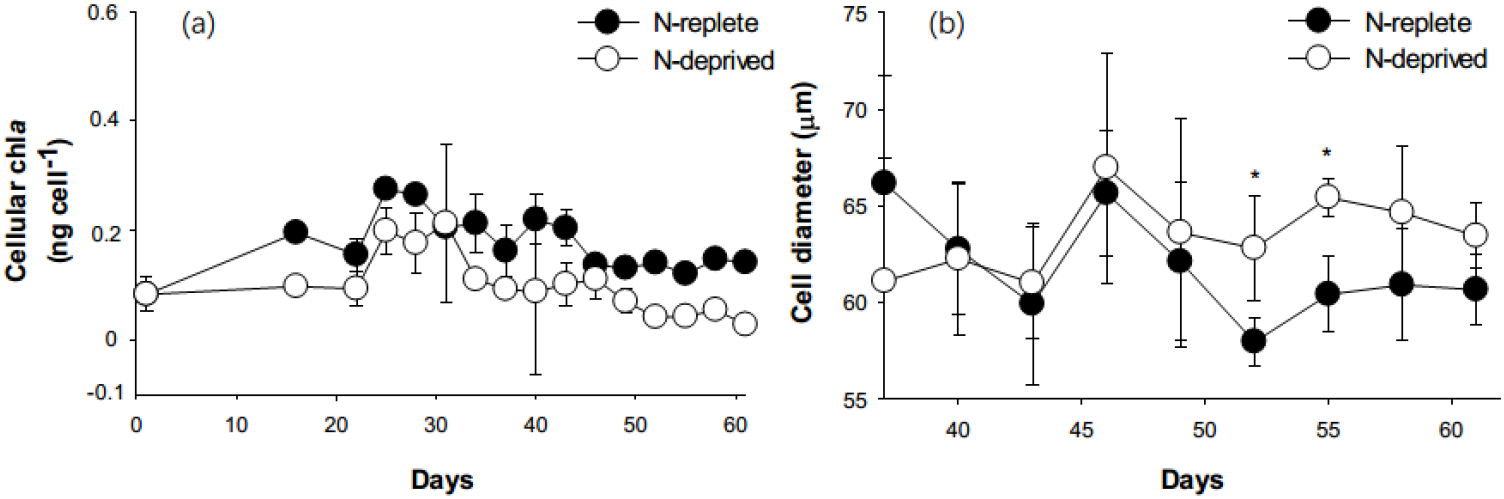
Chlorophyll *a* content (a) and cell size (b) of *G. balechii* under N-replete and N-limited conditions. Significant differences between the treatment groups are indicated by * (P < 0.05).

### QPCR quantification of 16S rDNA and CWH gene of *R. rosettiformans* strain GAMBA-01 under N-replete and N-limited conditions

We conducted qPCR for the 16S rDNA of *R. rosettiformans* strain GAMBA-01 to quantify copy numbers (proxy of abundance) in the early (day 40) and late (day 61) exponential growth phases of *G. balechii*. Considering equal DNA input in qPCR reactions, the relative abundance of the 16S rDNA was higher in the N-limited groups during both early and late exponential growth phases than in the N-replete cultures (Fig. 6). The copy number difference in the late exponential phase between the N-replete and N-limited cultures was significant (P < 0.05) (Fig. 6).

**FIG. 6.**
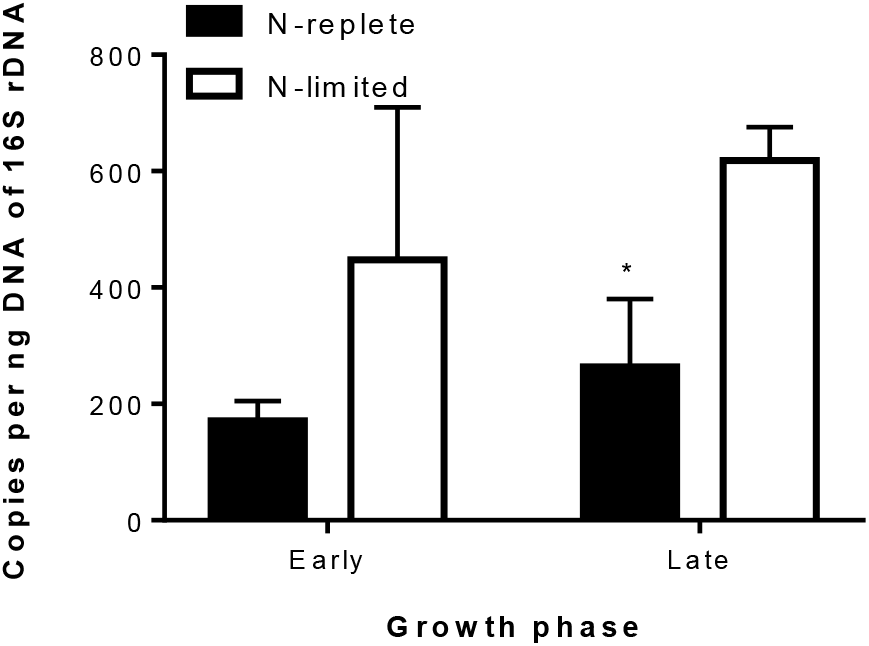
Abundance of *R. rosettiformans* GAMBA-01 16S rDNA gene in different N treatment groups of *G. balechii* during the early and late exponential phases. Significant difference between the treatment groups is indicated by * (P < 0.05).

The copy numbers of the CWH gene on day 40 and day 61 in the N-replete and N-limited groups of *G. balechii* were also quantified. Similar to the 16S rDNA, the CWH gene was more abundant in the N-limited group than in the N-replete group, and the difference was significant during the late exponential phase (P < 0.05) (Fig. 7). In contrast, no significant difference in CWH copy number was detected in the N-replete group between the early and late exponential phases (Fig. 7).

**FIG. 7.**
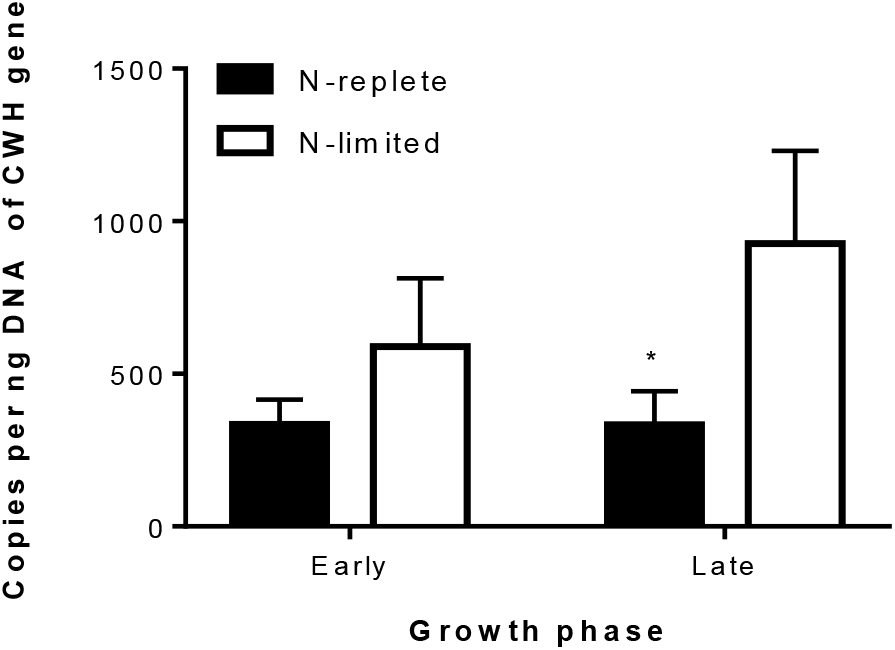
Abundance of *R. rosettiformans* GAMBA-01 CWH gene in different N treatment groups of *G. balechii* during the early and late exponential phases. Significant difference between the treatment groups is indicated by * (P < 0.05).

## DISCUSSION

Marine algae and bacteria, as primary producers and decomposers, are the dominant microorganisms in aquatic environments and their interactions play a major role in important biogeochemical processes [34, 35]. Numerous studies have reported how algae and the associated bacteria can benefit from their symbiotic relationships. For instance, algae can utilize vitamins generated by bacteria and meanwhile bacteria can utilize organic carbon produced by algal photosynthetic processes [2, 3, 7].

*Gambierdiscus* spp. are notorious benthic harmful algal bloom species mainly distributed in tropical and subtropical areas, commonly as an epiphyte of macro-algae or algal turf covering hard substrates in coral reef ecosystems [36]. The density of *Gambierdiscus* spp. in natural waters usually fluctuates widely within a short period; however, analyses of environmental factors such as temperature, salinity, or nutrients have not attributed these fluctuations to any of these factors [37]. It is possible that interactions with bacteria affect the population variation of *Gambierdiscus* spp. and alter the effect of the environmental factors. Previous studies have suggested that *Gambierdiscus* spp. are possibly mixotrophic microalgae and have the potential to assimilate and utilize some organic substances or nutrients released by bacteria [30, 36], which might explain why species in this genus can survive for more than 60 days when nutrients are scarce as observed in the present study.

However, to date, no symbiont bacterium for this dinoflagellate has been reported. In the present study, we isolated a strain of *Rhizobium* bacterium from the toxic benthic dinoflagellate *G. balechii* culture that harbors a CWH gene. A range of analyses using different methods, including physiological and molecular techniques, were used to identify and characterize this bacterial strain on *G. balechii*. As the first report of a *Rhizobium* bacterium associated with *G. balechii*, the study also provides a new model for studying the algae-bacteria associations.

### A *Rhizobium* bacterium from *G. balechii* culture

*Rhizobium*, a genus in the order Rhizobiales, is widely reported to be associated with higher plants [38]. Rhizobiales can elicit the formation of nodules on the roots or stems of their leguminous host to convert atmospheric N_2_ into ammonia using the enzyme nitrogenase, and then provide organic nitrogenous compounds such as glutamine or ureides to the host plants [38]. These plants, in return, provide the bacteria with organic compounds made via photosynthesis [38].

Considering that N sources can be limiting in the marine environment, particularly in the tropical coral reef ecosystems usually inhabited by *Gambierdiscus* species, associations of this alga with Rhizobiales bacteria are of particular interest, as their potential to fix N_2_ might offer advantages for the algae to thrive under the N-limited condition. Rhizobiales bacteria have been documented to exist in association with dinoflagellates, including the toxic *Prorocentrum lima* [39], *Alexandrium lusitanicum* [39], and *Alexandrium minutum* [40]. For example, *Hoeflea phototrophica* has been isolated from the cultures of *P. lima* and *A. lusitanicum* [39], and a species from the same bacterial genus (*H. alexandrii*) was isolated from *A. minutum* [40].

The Rhizobiales bacterium isolated from *G. balechii* culture was found to share a 99.93% identity with *R. rosettiformans* strain W3, which was isolated from groundwater in Lucknow, India [31]. This is the first report of the association of a Rhizobiales species with *Gambierdiscus*, as *Rhizobium* species have been documented in symbiotic relationships with green algae [24, 41] and brown algae [25, 42] in the marine ecosystem. The benefit of the association with *G. balechii* is unclear and can only be investigated using the information available regarding the related taxa. It has been reported that associations with Rhizobiales bacteria might play a crucial role in the fitness of *Alexandrium* strains, as species of the order Rhizobiales have been frequently reported during the blooms of *Alexandrium* spp. [43–45]. In addition, bacteria from this order have been found in the benthic assemblage of a coral reef ecosystem [46]. For instance, Rhizobiales bacteria were detected in the symbiotic microbiome of corals but not detected in the external coral colony of microbial community [47]. The bacterium from the genus *Parvibaculum* order Rhizobiales, was also detected on the surface of the symbiont dinoflagellate *Breviolum minutum* (formerly *Symbiodinium minutum*) [48] using scanning electron microscopy [49]. It is interesting that Rhizobiales have been frequently reported to be associated with benthic dinoflagellates from coral reef systems, such as *P. lima*, *B. minutum*, and now *G. balechii* documented in the present study. It is likely that these bacteria may benefit from the association with dinoflagellates and also promote the growth of benthic dinoflagellates, which is crucial for the coral community.

### Increased abundance of *R. rosettiformans* GAMBA-01 under N limitation and in the late exponential growth phase

Similar to previous studies on other species of dinoflagellates [50–52], N limitation decreased *G. balechii* growth rate and cell yield in the present study. Cellular chlorophyll *a* content, an important indicator of the photosynthetic process [53, 54], was reduced under N-nutrient stress. Chlorophyll *a* content also decreased during the late exponential phase in both N treatments, a growth stage when nutrient reduction to a limiting level is often expected. This suggests that N limitation reduces photosynthesis in *G. balechii*. As nutrient stress and aging have been shown to induce the production of extracellular polycarbohydrate substance [52, 55], the late stationary growth stage and N-limited culture condition are likely to create a favorable organic carbon environment for *Rhizobium* and other bacteria in the *G. balechii* cultures. Indeed, our results of qPCR analysis showed that *R. rosettiformans* GAMBA-01 (based on 16S rDNA and CWH) was more abundant in the N-limited cultures of *G. balechii* than in N-replete cultures; the difference was statistically significant during the late exponential phase when growth of *G. balechii* in the N-limited treatment was remarkably lower than in the N-replete treatment. This suggested that this bacterium may probably benefit from the association with *G. balechii* by obtaining organic matter such as carbohydrate released from the dinoflagellate, especially during the late exponential phase. In return, the dinoflagellate might survive at lower cell concentrations by obtaining the nutrients after ingesting the bacterium when N is limiting in the culture. Further investigations are required to verify this in the future.

### *bch*X as a paralog of *nif*H in *R. rosettiformans* GAMBA-01

As a species of *Rhizobium*, a known N_2_-fixing microorganism [56], we suspected that *R. rosettiformans* isolated from the *G. balechii* culture might be able to fix N_2_. N_2_ fixation is believed to be the evolutionary driver of many symbiotic systems, such as diatom-Richelia [57] and UCYN-A–haptophyte systems [58], in addition to the classical legume-*Rhizobium* system [23]. N_2_ fixation is mediated by the nitrogenous enzyme complex including the dinitrogenase reductase encoded by the *nif*H gene, which is considered one of the most genetically conserved genes [59, 60]. Using *nif*H universal primers, we attempted to detect and isolate this gene in our *R. rosettiformans* isolate. Sequence analysis of the PCR product obtained indicated that it was not a classical N_2_-fixing *nif*H gene. The *nif*H gene sequences are phylogenetically classified into five clusters [59, 61, 62]. Cluster I, composed almost entirely of sequences from the canonical FeMo nitrogenase [59, 60], consists of sequences from aerobic N_2_ fixers of Proteobacteria, Cyanobacteria, and Frankia [59, 60]. Cluster II is generally considered to be the alternative nitrogenase cluster as it contains sequences from FeFe and FeV nitrogenases, which differ from the conventional FeMo cofactor-containing nitrogenase [59]. Cluster III consists of anaerobic N fixers from bacteria and archaea including *Desulfovibrionaceae*, *Clostridia*, Spirochataes, and Methanobacteria [59]. Cluster IV and cluster V contain paralogs of *nif*H that are not involved in N_2_ fixation [59, 61]. Our sequence was a member of cluster V. The cluster V nitrogenase homologs, pigment biosynthesis complexes protochlorophyllide reductase, and chlorophyllide reductase, are not only homologous but are functionally analogous to nitrogenase, coupling ATP hydrolysis-driven electron transfer to substrate reduction [61]. As with nitrogenase, electrons flow from a NifH-like ATPase (BchL and BchX) to a NifDK-like putative heterotetramer, where the tetrapyrrole is bound (BchNB and BchYZ) [61].

*bch*X is responsible for the biosynthesis of bacteriochlorophyll in photosynthetic bacteria [26, 63], indicating the photosynthetic potential of *R. rosettiformans* GAMBA-01. The Rhizobiales bacterium *H. phototrophica* isolated from the benthic dinoflagellate *P. lima* culture is known to perform photosynthesis [39]. However, *H. phototrphica* was not able to fix N_2_ in the dinoflagellate culture [39]. Similarly, Rhizobiales bacteria were detected in the microbial community of *Alexandrium* spp.; however, N_2_-fixing activity was not detected and *nif*H could not be amplified under laboratory conditions [43]. Furthermore, a non-symbiotic *H. alexandrii* was isolated from the toxic dinoflagellate *A. minutum* AL1V [40]. These results together indicated that the Rhizobiales bacteria isolated from the *G. balechii* and other dinoflagellates are unlikely to be able to fix N_2_. More studies are required to elucidate the functions of *bch*X in these bacteria in dinoflagellate cultures.

### CWH gene and its potential functions

CWHs are a group of bacterial enzymes that are capable of hydrolyzing the stress-bearing peptidoglycan layer of their own cell wall [64, 65]. Some of these hydrolases can trigger cell lysis; therefore, they can be called autolysins or suicide enzymes [66]. Autolysins have been observed to be involved in a series of important cellular processes, including cell wall turnover, cell separation, competence, and flagellation (motility) [67].

The CWH gene isolated in the present study contains a conserved domain, *cwl*J, which has been reported to be involved in spore germination [32, 33]. Usually, spores of bacteria can be dormant and extremely resistant to environmental stress [68]. As a consequence, spores can survive for long periods in the absence of nutrients, whereas they can rapidly return to life (germination) once nutrients are available in their environment [69]. Germination requires degradation of the peptidoglycan layer termed the spore cortex, which can be performed by *cwl*J [32, 33]. Thus, *cwl*J is critical for spore germination in the presence of nutrients in the bacterial environment.

To our knowledge, CWH has not been studied previously in any algae-associated bacteria, and its function in *R. rosettiformans* GAMBA-01 remains unclear. It is possible that GAMBA-01 cells stay in the form of spores in the dinoflagellate culture during a specific phase to survive nutrient scarcity. Once nutrients return to the culture, CWH can be expressed to regulate spore germination of GAMBA-01. In the present study, we observed higher copy numbers of CWH in the N-limited cultures than in the N-replete cultures, especially in the late exponential phase. This indicated that N limitation of *G. balechii* can promote the growth of the GAMBA-01 strain, probably because this bacterium assimilates and utilizes organic matter released from *G. balechii*, which increases under N limitation, especially during the late exponential phase [52, 55]. Whether CWH functions in utilizing organic matter or in cell wall turnover remains an open question. One drawback of our study is that we did not assess the expression of this gene under contrasting N conditions or examine its photosynthetic ability, which should be included in the future investigations.

## CONCLUSION

Using bacteriological and molecular methods, we isolated a *Rhizobium* bacterium from the culture of the ciguatoxin-producing dinoflagellate *G. balechii*. This is the first report of the association of this bacterium with a *Gambierdiscus* species and of a *Rhizobium* species with any dinoflagellate. We found that this bacterium possesses a CWH-encoding gene, which might be related to spore germination, organic utilization, or cell wall turnover of this bacterium. As bacteria belonging to Rhizobiales typically are N_2_ fixers, we attempted to isolate *nif*H, which is commonly used as a marker gene for N_2_-fixing organisms. PCR with universal *nif*H primers did not amplify any authentic *nif*H gene; instead, we obtained a *nif*H homolog, *bch*X, which is known to exist in bacteria with photosynthetic ability. We observed an increase in the abundance of this bacterium as well as in CWH in the N-limited cultures of *G. balechii*, especially during the late exponential phase. While it is unlikely that this bacterium can perform N_2_ fixation in the *G. balechii* cultures, it is apparent that it can benefit from the co-culture by incorporating and utilizing the organic matter released from the dinoflagellate. On the other hand, *G. balechii* might be able to benefit from the association by using the bacterium as food or as a source of recycled nutritional substances for survival under low N-nutrient conditions. Further investigations are required to understand the roles of CWH and *bch*X and the functional interactions of the *Rhizobium* bacterium and *G. balechii*.

## MATERIALS AND METHODS

### Algal culture

*Gambierdiscus balechii* strain M1M10 was originally isolated from Kiribati in 2012 and was maintained in the State Key Laboratory of Marine Pollution, City University of Hong Kong. The culture was grown in autoclaved and 0.22-µm filtered artificial seawater with L1 medium nutrients (without silicate) [70]. The cultures were maintained at 25°C under a 12:12 light: dark regime at a photon flux of 100 μmol m^−2^s^−1^.

### Isolation of CWH-containing bacteria

A sample (10 mL) was removed from the *Gambierdiscus* culture and filtered through 3-μm membrane (Merck Millipore, Darmstadt, Germany). One hundred microliters of the filtrate, which was expected to contain bacteria, was spread onto a marine agar plate (Luria Bertani (LB) medium prepared using seawater and containing 1.5% agar). The plate was grown at 25 °C, under a 12:12 light: dark regime at a photon flux of 100 μmol m^−2^ s^−1^. Two days later, when colonies appeared, PCR was performed using colonies picked on a pipette tip directly as templates. To screen for bacteria that contained CWH (approximately 670 bp), common CWH gene primers (Table 1) were designed from conserved regions of this gene identified from an alignment of CWH sequences from various bacterial species. PCR amplification with these primers was performed under the following conditions: initial denaturation at 95 °C for 2 min, followed by 40 cycles of denaturation at 95 °C for 30 s, annealing at 55 °C for 30 s, and extension at 72 °C for 40 s, with a final extension cycle at 72 °C for 6 min. The PCR-positive bacterial clones were isolated into monospecific cultures for further analyses.

**Table 1.**
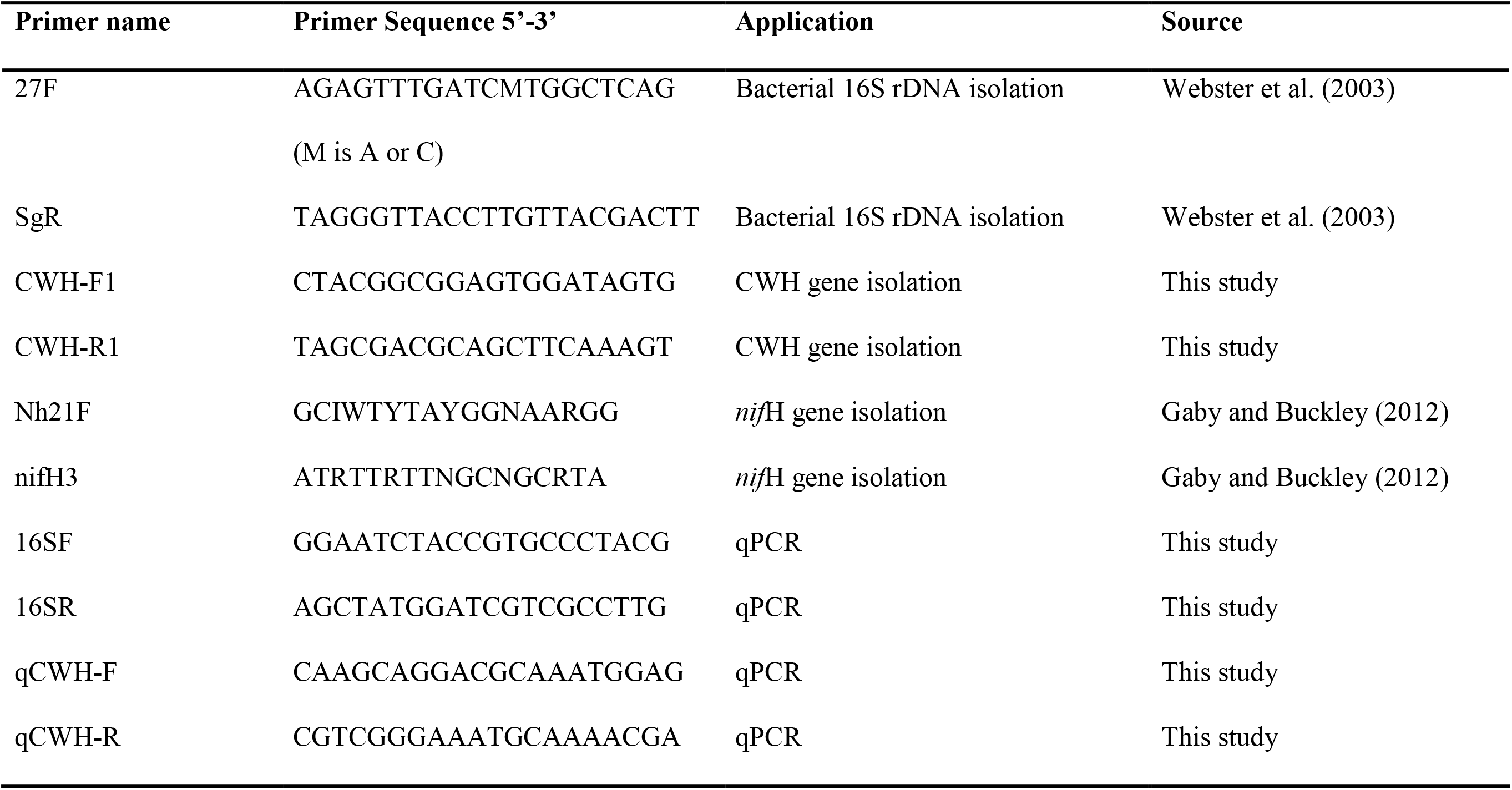
PCR primers used in the present study.

### DNA extraction, amplification, and sequencing of the 16S rDNA and CWH gene

Bacterial colonies that yielded positive results in CWH-PCR were selected and cultured in 50 mL Zobell 2216E broth at 25 °C for 24 h with shaking at 200 rpm. Bacterial cells were harvested via centrifugation at 3,901 g for 10 min and then transferred into a 2-mL micro-centrifuge tube with 0.5 mL DNA lysis buffer [19]. The samples were incubated at 55 °C for one day. DNA extraction was conducted using a CTAB protocol combined with the Zymo DNA clean and concentrator kit (Zymo Research Corp., Orange, CA, USA) as reported previously [71]. The 16S rDNA was amplified using the following parameters: 95 °C for 2 min, followed by 35 cycles of denaturation at 95 °C for 30 s, annealing at 56 °C for 30 s, and extension at 72 °C for 1 min, with a final extension cycle at 72 °C for 6 min. Primers 27F and SgR were used for the amplification of a 1.5-kb fragment of the 16S rDNA (Table 1) [19, 72]. The amplicons were subjected to Sanger sequencing (BGI, Shenzhen, China) directly.

### Amplification of potential *nif*H gene

Based on the 16S rDNA sequence, one of the bacterial isolates annotated as *Rhizobium* (see Results) was selected for further characterization. To determine whether this bacterium possesses the *nif*H gene, a *nif*H universal primer set (Nh21F and nifH3; Table 1) was used as described before [59] to amplify a 476-bp fragment. Amplification was performed using the following program: initial denaturation at 95 °C for 2 min, followed by 40 cycles of denaturation at 95 °C for 30 s, annealing at 41 °C for 30 s, and extension at 72 °C for 40 s, with a final extension cycle at 72 °C for 6 min. The amplicons were gel purified, cloned into the PMD-19T vector (Takara Biotechnology Co., Ltd., Beijing, China), and transformed into *Escherichia coli* DH5α competent cells (Takara Biotechnology Co., Ltd., Beijing, China). The resulting clones were randomly selected for Sanger sequencing.

### Nitrogen experiment

To grow *G. balechii* with coexisting bacteria, L1 medium (without silicate) was prepared with two different nitrate concentrations using autoclaved artificial seawater. For the N-limited cultures, the L1 medium was prepared with reduced nitrate concentration (4.41 µM). No more nitrogeneous compound was added during the experiment. N-replete cultures were set up as the control, in which 882 µM NaNO_3_ was added. Both control and N-limited treatments were performed in triplicate. Eight hundred milliliters xenic *G. balechii* was inoculated into the two sets of cultures in 1000-mL flasks. From day 1, samples were collected after every two days from each of the cultures for cell enumeration and measurement of cell size and chlorophyll *a* content. Algal cells were fixed in Lugol’ s solution and examined microscopically with a Sedgewick-Rafter counting chamber [73]. Daily population growth rate was calculated as µ= ln(*N*_*2*_/*N*_*1*_)/(*t*_*2*_-*t*_*1*_), where *N*_*2*_ and *N*_*1*_ are cell concentrations on day *t*_*2*_ and day *t*_*1*_, respectively. The mean cell size was measured as equivalent spherical diameter under the Nikon ECLIPSE 90i digital microscope (Nikon, Japan). Chlorophyll *a* content was determined using a microplate reader (BMG Polarstar Optima) after filtration of the sub-samples (5 to 10 mL) through Whatman GF/F glass circles (25 mm) (Merck KGaA, Darmstadt, Germany) and 90% acetone extraction of the filtrate for 24 h in the dark at 4 °C [74].

On day 40 (early exponential phase) and day 61 (late exponential phase), 10 mL of the *G. balechii* culture was collected from each treatment and was filtered using a 0.22-μm nitrocellulose membrane (Merck Millipore, Darmstadt, Germany). The membrane was then transferred into a 2-mL micro-centrifuge tube containing 1 mL DNA lysis buffer (containing 0.1 M EDTA, 1% sodium dodecyl sulfate, and 8 µg lysozyme). The samples were incubated at 55 °C for one week, with the prolonged incubation intended to maximize algal cell breakage, which has proven to be challenging in our laboratory. DNA extraction was conducted as described above.

### Quantitative PCR (qPCR) for the 16S rDNA and CWH-encoding gene of the target bacterium

Copy numbers of the 16S rDNA and CWH-encoding gene were determined using qPCR on the StepOnePlus real-time PCR system (Applied Biosystems, USA). Specific primers (16SF and 16SR; Table 1) were designed based on the 16S rDNA sequences obtained from the *Rhizobium* bacterium isolated in the present study. For the CWH gene, primers qCWH-F and qCWH-R were also designed based on the sequence obtained from our Sanger sequencing result (Table 1). The PCR products were purified and prepared in serial 10-fold dilutions to be used as standards. The same molar quantity of genomic DNA (0.01 ng) was used as the template to amplify the target gene from each group (N-replete and N-limited). Their copy numbers were calculated based on the standards, which were amplified on the same PCR runs as the samples. Each PCR was performed in a total volume of 10 µL containing 5 µL of 2 × iQSYBR Green supermix (Bio-Rad, USA), 200 nM of each primer, and 4 µL of 0.01 ng/µL DNA. The PCR program consisted of a denaturation step of 95 °C for 30 s, followed by 40 cycles of 95 °C for 5 s and 59 °C for 30 s. Each reaction had two technical replicates. Finally, all the PCR products were subjected to melting curve analysis to confirm primer specificity.

### Phylogenetic analysis

Phylogenetic trees were constructed to determine the affinity of the bacterium strain to the documented taxa of bacteria. 16S rDNA sequences as well as amino acid sequences closely related to the CWH homolog isolated in this study were obtained from National Center of Biotechnology Information (NCBI) to combine with the sequences generated from this study as a dataset. The alignment of these 16S rDNA sequences was performed using ClustalW via MEGA7 [75]. MUSCLE via MEGA7 was employed for the analysis of CWH amino acid sequences. Prior to maximum likelihood (ML) phylogenetic analysis, the best DNA substitution model fitting for 16S rDNA sequences was T92+G which was selected using the Akaike information criterion [76]. The best amino acid substation model fitting for CWH amino acid sequences was WAG+G [76]. ML phylogenetic trees were created using MEGA7 with 1,000 bootstraps [75]. Neighbor-Joining analysis conducted using MEGA7 [75] yielded similar tree topology.

### Statistical analysis

To compare the differences in the variables between the N-replete and N-limited groups, analysis of variance (ANOVA) was conducted using the SPSS Statistics 17.0 software package. Significant difference was set at P < 0.05. All data presented are means with standard deviation calculated from the triplicate cultures under each N condition.

## ACKNOWLEDGEMENTS

We thank Mr. Zhang Kaidian, Dr. Zhang Hua, and Dr. Shi Xinguo for their assistance in this study. The study was supported by the Collaborative Research Fund from the Research Grant Council [C1012-15G] of Hong Kong.

## CONFLICT OF INTEREST

The authors declared that they have no conflicts of interest to this work.

## REFERENCES

1. Cole, J.J., Interactions between bacteria and algae in aquatic ecosystems. Annual Review of Ecology Systematics, 1982. 13(1): p. 291–314.

2. Tang, Y.Z., F. Koch, and C.J. Gobler, Most harmful algal bloom species are vitamin B1 and B12 auxotrophs. Proceedings of the National Academy of Sciences, 2010. 107(48): p. 20756–20761.

3. Croft, M.T., et al., Algae acquire vitamin B 12 through a symbiotic relationship with bacteria. Nature, 2005. 438(7064): p. 90–93.

4. Menzel, D.W. and J.P. Spaeth, Occurrence of vitamin b12 in the sargasso sea. Limnology and Oceanography, 1962. 7(2): p. 151–154.

5. Haines, K.C. and R.R. Guillard, Growth of vitamin b12-requiring marine diatoms in mixed laboratory cultures with vitamin b12-producing marine bacteria. Journal of Phycology, 1974. 10(3): p. 245–252.

6. Swift, D.G. and R.R. Guillard, Unexpected response to vitamin b12 of dominant centric diatoms from the spring bloom in the gulf of maine (northeast atlantic ocean). Journal of Phycology, 1978. 14(4): p. 377–386.

7. Kuo, R.C. and S. Lin, Ectobiotic and endobiotic bacteria associated with Eutreptiella sp. isolated from Long Island Sound. Protist, 2013. 164(1): p. 60–74.

8. Golterman, H., Role of phytoplankton in detritus formation. Memorie dell’Istituto Italiano di Idrobiologia, 1972. 29: p. 89–103.

9. Bloesch, J., P. Stadelmann, and H. Bührer, Primary production, mineralization, and sedimentation in the euphotic zone of two Swiss lakes. Limnology and Oceanography, 1977. 22(3): p. 511–526.

10. Axler, R.P., G.W. Redfield, and C.R. Goldman, The importance of regenerated nitrogen to phytoplankton productivity to phytoplankton productivity in a subalpine lake. Ecology, 1981. 62(2): p. 345–354.

11. Nakanishi, K., et al., Bacteria that induce morphogenesis in Ulva pertusa (Chlorophyta) grown under axenic conditions. Journal of Phycology, 1996. 32(3): p. 479–482.

12. Delucca, R. and M.D. McCracken, Observations on interactions between naturally-collected bacteria and several species of algae. Hydrobiologia, 1977. 55(1): p. 71–75.

13. Amin, S., et al., Interaction and signalling between a cosmopolitan phytoplankton and associated bacteria. Nature, 2015. 522(7554): p. 98–101.

14. Jeong, H.J., et al., Feeding and grazing impact by small marine heterotrophic dinoflagellates on heterotrophic bacteria. Journal of Eukaryotic Microbiology, 2008. 55(4): p. 271–288.

15. Alavi, M., et al., Bacterial community associated with Pfiesteria-like dinoflagellate cultures. Environmental Microbiology, 2001. 3(6): p. 380–396.

16. Anderson, J.M. and A. Macfadyen. The role of terrestrial and aquatic organisms in decomposition processes. in Proceedings of the 17th Symposium of the BES. 1976.

17. Sule, P. and R. Belas, A novel inducer of Roseobacter motility is also a disruptor of algal symbiosis. Journal of bacteriology, 2013. 195(4): p. 637–646.

18. Alonso, C., et al., High local and global diversity of Flavobacteria in marine plankton. Environmental microbiology, 2007. 9(5): p. 1253–1266.

19. Wang, C., et al., Glyphosate shapes a dinoflagellate-associated bacterial community while supporting algal growth as sole phosphorus source. Frontiers in microbiology, 2017. 8: p. 2530.

20. Wang, X., et al., Lysis of a red-tide causing alga, Alexandrium tamarense, caused by bacteria from its phycosphere. Biological Control, 2010. 52(2): p. 123–130.

21. Biebl, H., et al., Dinoroseobacter shibae gen. nov., sp. nov., a new aerobic phototrophic bacterium isolated from dinoflagellates. International journal of systematic evolutionary microbiology, 2005. 55(3): p. 1089–1096.

22. Amin, S.A., et al., Photolysis of iron–siderophore chelates promotes bacterial– algal mutualism. Proceedings of the National Academy of Sciences, 2009. 106(40): p. 17071–17076.

23. Mitsustin, E. and V. Sil’nikova, Biological fixation of atmospheric nitrogen. 1968, Moscow. 531.

24. Kim, B.-H., et al., Role of Rhizobium, a plant growth promoting bacterium, in enhancing algal biomass through mutualistic interaction. Biomass and Bioenergy, 2014. 69(2014): p. 95–105.

25. Dittami, S.M., et al., Genome and metabolic network of “Candidatus Phaeomarinobacter ectocarpi” Ec32, a new candidate genus of Alphaproteobacteria frequently associated with brown algae. Frontiers in genetics, 2014. 5: p. 241.

26. Nomata, J., et al., A second nitrogenase-like enzyme for bacteriochlorophyll biosynthesis reconstitution of chlorophyllide a reductase with purified x-protein (bchX) and yz-protein (bchY-bchZ) from Rhodobacter capsulatus. Journal of Biological Chemistry, 2006. 281(21): p. 15021–15028.

27. Burke, D.H., J.E. Hearst, and A. Sidow, Early evolution of photosynthesis: clues from nitrogenase and chlorophyll iron proteins. Proceedings of the National Academy of Sciences, 1993. 90(15): p. 7134–7138.

28. Dai, X., et al., Taxonomic assignment of the benthic toxigenic dinoflagellate Gambierdiscus sp. type 6 as Gambierdiscus balechii (Dinophyceae), including its distribution and ciguatoxicity. Harmful Algae, 2017. 67: p. 107–118.

29. Lartigue, J., et al., Nitrogen source effects on the growth and toxicity of two strains of the ciguatera-causing dinoflagellate Gambierdiscus toxicus. Harmful Algae, 2009. 8(5): p. 781–791.

30. Price, D.C., et al., Analysis of Gambierdiscus transcriptome data supports ancient origins of mixotrophic pathways in dinoflagellates. Environmental microbiology, 2016. 18(12): p. 4501–4510.

31. Kaur, J., M. Verma, and R. Lal, Rhizobium rosettiformans sp. nov., isolated from a hexachlorocyclohexane dump site, and reclassification of Blastobacter aggregatus Hirsch and Müller 1986 as Rhizobium aggregatum comb. nov. International journal of systematic and evolutionary microbiology, 2011. 61(5): p. 1218–1225.

32. Heffron, J.D., B. Orsburn, and D.L. Popham, Roles of germination-specific lytic enzymes CwlJ and SleB in Bacillus anthracis. Journal of bacteriology, 2009. 191(7): p. 2237–2247.

33. Li, Y., et al., Activity and regulation of various forms of CwlJ, SleB, and YpeB proteins in degrading cortex peptidoglycan of spores of Bacillus species in vitro and during spore germination. Journal of bacteriology, 2013. 195(11): p. 2530–2540.

34. Simon, N., et al., Kinetics of attachment of potentially toxic bacteria to Alexandrium tamarense. Aquatic microbial ecology, 2002. 28(3): p. 249–256.

35. Danger, M., et al., Control of phytoplankton–bacteria interactions by stoichiometric constraints. Oikos, 2007. 116(7): p. 1079–1086.

36. Sakami, T., et al., Effects of epiphytic bacteria on the growth of the toxic dinoflagellate Gambierdiscus toxicus (Dinophyceae). Journal of experimental marine biology and ecology, 1999. 233(2): p. 231–246.

37. Shmukler, Y.B. and D.A. Nikishin, Ladder-shaped ion channel ligands: Current state of knowledge. Marine drugs, 2017. 15(7): p. 232.

38. Fisher, R.F. and S.R. Long, Rhizobium–plant signal exchange. Nature, 1992. 357(6380): p. 655–660.

39. Biebl, H., et al., Hoeflea phototrophica sp. nov., a novel marine aerobic alphaproteobacterium that forms bacteriochlorophyll a. International journal of systematic evolutionary microbiology, 2006. 56(4): p. 821–826.

40. Palacios, L., et al., Hoeflea alexandrii sp. nov., isolated from the toxic dinoflagellate Alexandrium minutum AL1V. International Journal of Systematic and Evolutionary Microbiology, 2006. 56(8): p. 1991–1995.

41. Hodkinson, B.P. and F. Lutzoni, A microbiotic survey of lichen-associated bacteria reveals a new lineage from the Rhizobiales. Symbiosis, 2009. 49(3): p. 163–180.

42. Dittami, S.M., et al., Host–microbe interactions as a driver of acclimation to salinity gradients in brown algal cultures. The ISME journal, 2016. 10(1): p. 51–63.

43. Wiese, M., Investigations into abiotic and biotic factors regulating saxitoxin synthesis in the dinoflagellate genus Alexandrium. 2012, University of New South Wales: Australia.

44. Green, D.H., et al., Phylogenetic and functional diversity of the cultivable bacterial community associated with the paralytic shellfish poisoning dinoflagellate Gymnodinium catenatum. FEMS Microbiology Ecology, 2004. 47(3): p. 345–357.

45. Zhou, J., et al., Microbial community structure and associations during a marine dinoflagellate bloom. Frontiers in microbiology, 2018. 9.

46. Barott, K.L., et al., Microbial diversity associated with four functional groups of benthic reef algae and the reef-building coral Montastraea annularis. Environmental microbiology, 2011. 13(5): p. 1192–1204.

47. Ainsworth, T.D., et al., The coral core microbiome identifies rare bacterial taxa as ubiquitous endosymbionts. The ISME journal, 2015. 9(10): p. 2261–2274.

48. LaJeunesse, T.C., et al., Systematic revision of Symbiodiniaceae highlights the antiquity and diversity of coral endosymbionts. Current Biology, 2018. 28(16): p. 2570–2580. e6.

49. Shoguchi, E., et al., Draft assembly of the Symbiodinium minutum nuclear genome reveals dinoflagellate gene structure. Current biology, 2013. 23(15): p. 1399–1408.

50. Zhang, C., et al., Suppression subtraction hybridization analysis revealed regulation of some cell cycle and toxin genes in Alexandrium catenella by phosphate limitation. Harmful Algae, 2014. 39: p. 26–39.

51. Li, M., et al., Phosphorus deficiency inhibits cell division but not growth in the dinoflagellate Amphidinium carterae. Frontiers in microbiology, 2016. 7: p. 826.

52. Vidyarathna, N.K. and E. Granéli, Physiological responses of Ostreopsis ovata to changes in N and P availability and temperature increase. Harmful algae, 2013. 21: p. 54–63.

53. Krause, G. and E. Weis, Chlorophyll fluorescence and photosynthesis: the basics. Annual review of plant biology, 1991. 42(1): p. 313–349.

54. Platt, T., D.S. Rao, and B. Irwin, Photosynthesis of picoplankton in the oligotrophic ocean. Nature, 1983. 301(5902): p. 702.

55. Vanucci, S., et al., Effects of different levels of N-and P-deficiency on cell yield, okadaic acid, DTX-1, protein and carbohydrate dynamics in the benthic dinoflagellate Prorocentrum lima. Harmful Algae, 2010. 9(6): p. 590–599.

56. Carvalho, F.M., et al., Genomic and evolutionary comparisons of diazotrophic and pathogenic bacteria of the order Rhizobiales. BMC microbiology, 2010. 10(1): p. 37.

57. Foster, R.A. and J.P. Zehr, Characterization of diatom–cyanobacteria symbioses on the basis of nifH, hetR and 16S rRNA sequences. Environmental Microbiology, 2006. 8(11): p. 1913–1925.

58. Krupke, A., et al., The effect of nutrients on carbon and nitrogen fixation by the UCYN-A–haptophyte symbiosis. The ISME journal, 2015. 9(7): p. 1635.

59. Gaby, J.C. and D.H. Buckley, A comprehensive evaluation of PCR primers to amplify the nifH gene of nitrogenase. PloS One, 2012. 7(7): p. e42149.

60. Zehr, J.P., M.T. Mellon, and S. Zani, New nitrogen-fixing microorganisms detected in oligotrophic oceans by amplification of nitrogenase (nifH) genes. Applied and Environmental Microbiology, 1998. 64(9): p. 3444–3450.

61. Raymond, J., et al., The natural history of nitrogen fixation. Molecular biology and evolution, 2004. 21(3): p. 541–554.

62. Ininbergs, K., et al., Composition and diversity of nifH genes of nitrogen-fixing cyanobacteria associated with boreal forest feather mosses. New Phytologist, 2011. 192(2): p. 507–517.

63. Aagaard, J. and W. Sistrom, Control of synthesis of reaction center bacteriochlorophyll in photosynthetic bacteria. Photochemistry and photobiology, 1972. 15(2): p. 209–225.

64. Parisien, A., et al., Novel alternatives to antibiotics: bacteriophages, bacterial cell wall hydrolases, and antimicrobial peptides. Journal of applied microbiology, 2008. 104(1): p. 1–13.

65. Ishikawa, S., et al., Regulation of a new cell wall hydrolase gene, cwlF, which affects cell separation in Bacillus subtilis. Journal of Bacteriology, 1998. 180(9): p. 2549–2555.

66. Rogers, H.J., H.R. Perkins, and J.B. Ward, Microbial cell walls and membranes. Vol. 541. 1980: Springer.

67. Nugroho, F.A., et al., Characterization of a new sigma-K-dependent peptidoglycan hydrolase gene that plays a role in Bacillus subtilis mother cell lysis. Journal of Bacteriology, 1999. 181(20): p. 6230–6237.

68. Nicholson, W.L., et al., Resistance of Bacillus endospores to extreme terrestrial and extraterrestrial environments. Microbiology and Molecular Biology Reviews, 2000. 64(3): p. 548–572.

69. Setlow, P., Spores of Bacillus subtilis: their resistance to and killing by radiation, heat and chemicals. Journal of applied microbiology, 2006. 101(3): p. 514–525.

70. Guillard, R. and P. Hargraves, Stichochrysis immobilis is a diatom, not a chrysophyte. Phycologia, 1993. 32(3): p. 234–236.

71. Zhang, H., D. Bhattacharya, and S. Lin, Phylogeny of dinoflagellates based on mitochondrial cytochrome b and nuclear small subunit rdna sequence comparisons. Journal of Phycology, 2005. 41(2): p. 411–420.

72. Webster, G., et al., Assessment of bacterial community structure in the deep sub-seafloor biosphere by 16S rDNA-based techniques: a cautionary tale. Journal of Microbiological Methods, 2003. 55(1): p. 155–164.

73. Lin, X., et al., Alkaline phosphatase gene sequence characteristics and transcriptional regulation by phosphate limitation in Karenia brevis (Dinophyceae). Harmful Algae, 2012. 17: p. 14–24.

74. Ritchie, R.J., Consistent sets of spectrophotometric chlorophyll equations for acetone, methanol and ethanol solvents. Photosynthesis research, 2006. 89(1): p. 27–41.

75. Kumar, S., G. Stecher, and K. Tamura, MEGA7: molecular evolutionary genetics analysis version 7.0 for bigger datasets. Molecular biology and evolution, 2016. 33(7): p. 1870–1874.

76. Nei, M. and S. Kumar, Molecular evolution and phylogenetics. 2000: Oxford university press.

